# Proliferation of lung epithelial cells is regulated by the mechanisms of autophagy upon exposure of soots

**DOI:** 10.1101/2020.09.19.304725

**Authors:** Rituraj Niranjan, Kaushal Prasad Mishra, Ashwani Kumar Thakur

## Abstract

Soots are known to cause many diseases in humans but their underlying mechanisms of toxicity are still not known. Here, we report that, soots induce cell proliferation of lung epithelial cells via modulating autophagic pathways. Fullerene soot and diesel exhaust particles (DEP) induced cell proliferation of lung epithelial, A549 cells, via distinct autophagic mechanisms and did not cause cell death. Exposure of fullerene soot protected cell death of A549 cells, caused by hydrogen peroxide and inhibited LPS-induced autophagy. Fullerene soot co-localize with the autophagic proteins and inhibited starvation-induced autophagy (downregulated ATG-5, beclin-1, p62 and LC3 expressions) independent of its antioxidant properties. Similarly, it decreased expression profile of autophagic genes and upregulated proliferation responsive gene, Ki-67, in mice. We observed that, expressions of fullerene soot responsive genes (Beclin-1, ATG-5 and p62) were reverted by Akt Inhibitor X, indicating an important role of Akt pathway. On the other hand, DEP up-regulated expressions of autophagy genes. Akt Inhibitor X and Etoricoxib, did not attenuate DEP-induced cell proliferation and autophagic response. However, autophagic inhibiter 3-MA and chloroquine has significantly inhibited DEP-induced cell roliferation. In conclusion, distinct autophagic mechanisms are operational in cell proliferation of lung epithelial cells in response to soots and may have implication in diseases. The proliferation inducing potential of fullerene soot may be utilized as a regenerative agent in autophagy-associated disorders.

## Introduction

Ambient soot/black carbon and traffic related air pollution are responsible for the development of many diseases of humans and animals (Abas et al. 2004; Niranjan, Thakur 2017; Hedman et al. 2015; Jacquemin et al. ; Ji et al. 2015). Soot is consist of carbonaceous particles, free and fused polycyclic aromatic hydrocarbons (PAHs) which are formed after the incomplete combustion of gasoline, diesel and other petroleum fuels (Niranjan, Thakur 2017; Boffetta et al. 1997). Poly aromatic hydrocarbons (PAH) in soot are considered to be the main carcinogenic compound (Wang et al. 2001; Boffetta et al. 1997). Among many soots, diesel exhaust particles are considered as the main source of environmental soot and responsible for increase rate of morbidity and mortality in humans (Patel et al. 2013; Reynolds, Richards 2001). The carbon particles from wood smoke and road traffic also produce allergic adjuvant effects and exacerbate lung associated disorders (Samuelsen et al. 2008). Other model soots, such as fullerene soot (carbon black and nanocaged structure) also share same kind of toxicological responses with environmental soot. However, the role of both types of soots in causing biological effects on respiratory system are not extensively investigated (Harhaji et al. 2007; Baierl et al. 1996). Nevertheless, some mechanisms related to biological effects are known for them. For example, environmental soot (like diesel exhausts particles), significantly induces reactive oxygen species in the lungs of humans along with the experimental animals (Hussain et al. 2009; Hussain et al. 2010). On the other hand, reactive oxygen species (ROS) generation can lead to autophagy of cells. (Zhou et al. 2011; Altman, Rathmell 2009) (Duan et al. 2014) It is also widely accepted that, nanoparticles induces autophagy in a variety of models by more than one mechanisms (Afeseh Ngwa et al. 2011). Notably, soot also exist in nanoparticles forms and therefore may affect autophagic mechanisms by various ways (Andon, Fadeel 2013). Considerable studies have described that, a defect in autophagy is connected to development of lung pathologies, however, the underlying mechanisms are still unknown (Aggarwal et al. 2016). Thus to understand the biological effect and fundmental mechanism of soot on lungs, the presnt study was designed, where in fullerene soot and disesel soot were used first on lung epithelial cells to know the biological effect on cell viability and later the observations were assessed on in vivo system to investigate the effect of soot on lungs tissue. We found that in presence of fullerene soot, lung epeithelial cells show proliferation by reducing autophagy. This effect was found to be dependent on the Akt pathway. The same effect was observed in mouse lung cells where tissue proliferation and underexpression of autophagic genes were evident. Thus we have shown that soot induced cell proliferation can cause a hyperplasia-like tissue development which may in turn lead to tissue remodeling and lung diseases. Our work thus projects a glimpse into the consequences of soot exposure to human lungs.

## Materials and methods

### Chemicals and Reagents

Fullerene soot from Sigma Aldrich (product number 572497), primary rabbit polyclonal antibodies against cleaved caspases-3 and light chain 3 (LC3) were procured from Cell Signaling Technology (Danvers, MA, USA). APG-5, Ki-67. Beclin-1, p53 and secondary anti-rabbit HRP conjugated antibodies were procured from the Santa Cruz Biotechnology (Santa Cruz, USA). Primary rabbit polyclonal anti-p62 antibodies and secondary anti-rabbit Alexa *Fluor*® *488* conjugated antibodies were purchased from the life technologies (Invitrogen, Carlsbad, CA, USA). All other chemicals were purchased from Sigma-Aldrich (St. Louis, MO, USA). Diesel exhaust particles were collected using the standard procedure as described before with desired modifications (Gangwar et al. 2011).

### Cell lines used

Human lung epithelial cell line A549, Human embryonic kidney HEK 293, Human neuroblastoma SH-SY5Y cells, were used in the study. All these cell lines were purchased from National Centre for Cell Sciences, Pune, India and since then continuously maintained.

### Culture and passaging of the cells

DMEM or DMEM-ham’s F12 nutrient mixture was used for culturing the cells. In this neutrient mixture medium, 10% heat-inactivated fetal bovine serum was added. The culture of the cells was maintained at 37°C in a humidified atmosphere of 5% CO_2_/95% air. The pH of the medium was adjusted to nearly 7.4 with the help of 0.1 N NaOH / 0.1 N HCl followed by filter sterilization by Merk-Millipore filters of pore size 0.22 μm. At first sterility of the medium was checked keeping medium in CO_2_ incubator for at least 48 hours. Subculture of the cells was done using trypsinization as per the requirements. Depending on the confluence cell suspension was distributed into 2-3 flasks and 4 ml fresh medium was added to each flask.

### Preparartion of suspension of Fulleelrene soot and DEP in medium

Fullerene soot was obtained from sigma and then a stock of the fullereane soot (4 mg in 2ml) was directly made in medium (DMEM: Ham’s F12). In a similar way, DEP was also dissolved directly in the medium. The further dilutions of the required concentrations (125 µg/ml to 2000 µg/ml) were made in the ame medium. The dissolution of the soot was enchanced by vortexing it to many times with regular intervals for a period of 1 hours after mixing. Exposure to the cells with different concentrations of soots were given inside biosafety cabinet various time periods as per the requirments of the experiments.

### Cell viability assay by MTT

MTT assay was performed to determine cell viability (Mosmann 1983). Cells (10, 000/ well) were seeded in 96-well plate. After exposure with different concentrations of soots or inhibitors (as metioned in indivisual results sections) MTT salt 3-(4,5-dimethylthiazol-2-yl)-2,5-diphenyltetrazolium bromide (20 μl/well containing 100 μl of cell suspension; 5 mg/ml in PBS) was added. Color was read at 530 nm, using multi-well microplate reader. For each group minimum 3 to 6 replicates were used.

### Cell proliferation assay using fluorescence based kit

Cell proliferation in culture was measured by the CyQUANT® cell proliferation assay kit as per the protocol described by the provider (Thermo Fisher scientific). In brief, 4000 cells per well were seeded in the 96 well plate and were treated with different soot concentrations (ranging from 50 to 2000 µg/ml). Cell proliferation was determined by using the kit procedure as per the manufacturer’s instructions.

### Measurement of ROS generation

Measurement of intracellular generation of reactive oxygen species (ROS) was accomplished using fluorescence dye dichloro-dihydro-fluorescein diacetate (DCF-DA). A549 cells were seeded at the cell density of 1×10^4^ cells/well in 96 well plates. Following the treatment with soot, or inducer LPS 50 µg/ml and soot 250 µg/ml), medium was aspirated and DCF-DA (10 µM final concentrations) in phenol red free HBSS buffer (100 µl/well) was added to the plate. A minimum incubation period of 30 minute was given in dark at 37°C in CO_2_ incubator. Fluorescence measurement was done at excitation of 485 nm and emission at 530 nm, by fluorescence reader (Perkin Elmer).

### Measurement of soot internalization using methanol treatment

The determination of internalization of soot was accomplished by methanol treatment of soot. Initially soot was weighed and mixed with the methanol and then it was vortexed for 1 hour intermittently for mixing. Fullerene soot was incubated with methanol for 24 hours and then it was coated in the 96 well plates having cover slips in their bottom. These plates were allowed to dry in the laminar air flow for a period until the extra methanol is evaporated and the plates have only fullerene soot. A549 cells were then plated on the cover slips and allowed for a period of 24 hours for the attachment. After the incubation period the cells were fixed with the paraformaldehyde at 4°C. internalization of fullerene soot was measured using fluorescence microscope. Internalized particles gives red fluorescence which were captured using fluorescenece microscope.

### Measurements of organic carbon and elemental carbon by using OCEC instrument

The measurement of organic carbon and inorganic carbon contents before and after the soot treatment was done using the semi-continuous OCEC, carbon aerosol analyzer, instrument as per the manufactures protocol (Sunset Laboratory, Inc). In brief, A549 cells (1×10^5^/ml) were exposed to fullerene soot (1000 µg/ml) for different time intervals (4 hour, 8 hours, 12 hours). Cells were then collected and lysed in lysis biffer containing 150 mM Nacl. The lysed cells were then centrifuged at 400g for removal of debris. Cell debris was discarded and supernatants were kept at the filters for analysis by a OCEC instrument. The experiments was conducted with duplicates and repeated once.

### Animal studies

8 to 12 weeks old BALB/c mice were used in this study. Initially mice were divided in two groups, namely control and soot (Fullerene soot) treated mice .Each group consisted of atleast 4 mice.Institutional animal ethical clearance was obtained to conduct the use of exprementation on the mice as per the Committee for the Purpose of Control and Supervision of Experiments on Animals (CPCSEA) guidelines. Mice were anesthetized using isoflurane and then intranasal challenged with fullerene soot. 100 µg of soot prepared in saline (by mixing with saline using vortex) (was given to per mice per dose in 50 µl of volume. Two intranasal doses were given to the mice in 12-day period, in a interval of 6 days. At last mice were sacrificed and organs were isolated. Tissues were fixed in the neutral buffered formalin and later were analyzed for different markers.

### Immunocytochemistry for protein localization studies

Six well plate containing cover slips at their bottom were used to seed the cells (2×10^5^/well). Treatment of either soots or inhibitors (concentration) was given after 24h. After the incubation period, cells were fixed with 4% para-formaldehyde. Subsequently, cells were treated with 0.05% H_2_O_2_ in methanol for one hour at room temperature in dark with mild agitation to make them permeable. Cells were blocked by blocking buffer (0.02% BSA + 0.002 % Triton-X 100) for a period of 30 minute at room temperature. After blocking, treatment with primary antibodies at a dilution of 1:100 was given to the cells with mild agitation for overnight at 4°C. Cells were then treated with secondary Alexaflour488-conjugated antibodies, at 1:200 dilutions in blocking buffer for one hour at room temperature with mild agitation. The cells were washed with PBS after each step. Images were captured by upright fluorescence microscope at different magnifications.

### Immunohistochemistry of the tissue sections

The tissues of mice were fixed in neutral buffered formalin, embedded in paraffin and were cut into 5 µM thin sections. Sections were fixed to positive charged slides. The endogenous peroxidase activity in the tissues was quenched using 0.3% hydrogen peroxide in methanol. Sections were blocked by nonspecific protein blocking with 1% BSA. Tissue sections were then incubated with different antibodies overnight at 4°C at 1:100 dilutions, followed by 1:200 dilutions of HRP-conjugated anti-rabbit IgG secondary antibody for 1 hour at room temperature. Subsequently these slides were developed using DAB-kit and further counterstained with nuclear fast red or hematoxylin. Negative controls include secondary antibody alone without the primary antibody.

### Western immunoblotting analysis for the expression of proteins

Cell lysates of A549 cells after incubation with different treatments (controls and exposed) were prepared in 300 µl of lysis buffer, containing, 50 mM Tris-HCl, 1 mM EDTA, 100 mM NaCl, 100 µg/ml phenylmethylsulfonyle fluoride (PMSF) and protein inhibitor cocktail. The obtained lysates were then centrifuged at 10,000 g for 5 minutes at 4°C. Protein estimation in supernatant was estimated using BCA kit provided by pierce (thermo fisher) company. The linear range values of protein estimation were used to calculate the exact concentrations using different dilutions. Samples were mixed with NuPAGE, 4× loading buffer containing 200 mM dithiothreitol (DTT). In most of the cases, 25 to 100 µg of protein was separated on 8-12 % SDS-PAGE, and subsequently transferred to the PVDF membranes for the particular analysis. The blocking of membrane was done by blocking buffer (5% non-fat dry milk, 100 mM NaCl, 10 mM Tris pH = 7.5, 0.1% tween-20) for one hour at room temperature. Standardization of signal was done using different concentration of protein or antibodies to avoid saturation effects and to get optimized results in linear range. Membranes were then treated with different protein specific primary antibodies (at 1:500 to 5000 dilutions) at 4°C for overnight. Membranes were washed 5 times with TBST. Membranes were then treated with HRP-conjugated secondary antibodies for one hour (from 1: 2000 to 1: 10000 dilutions) at room temperature. After that blots were developed by using ECL (Enhanced chemiluminescence) system provided by the pierce. Densitometry analyses of bands were accomplished by the gel documentation system (Bio Red, U.S.A.).

### Statistical analysis

Results, from the majority of experiments are expressed as mean ± S.E.M or otherwise specified. The accomplishment of statistical analysis was done by one-way analysis of variance (ANOVA), followed by Newman-Keuls test as post-hoc test. The p value <0.05 was considered statistically significant. Graph-Pad prism 5 software was adopted for the statistical analysis.

## Results

### Effect of fullerene soot on the cell morphology, cell proliferation and apoptosis of lung epithelial cells, A549

At first, we have tested effect of fullerene soot on lung airway epithelial cells, the first site of entry of soots to human body. In order to assess toxic effect of fullerene soot, we have tested its effect on the cell viability or cell proliferation of A549 cells. As seen in Fig. 1A, soot exposure did not cause cell death of these cells. Soot concentrations (250, 500, 1000, and 2000 µg/ml) have increased MTT score in 24 hours of incubation period of cells. As shown in Fig. 1 D, soot did not cause death of these cells. Morphologically, soot exposed cells are big in size and diffentiating than control cells. Notably, some particles of soot are seen to be internalized by the cells exposed to fullerene soot. In addition to MTT assay, we also tested fullerene soot-induced apoptosis of A549 cells. Apoptosis of A549 cells in response to soot was measured by DAPI nuclear staining. To achieve this, 1000 µg/ml of soot concentration was exposed to the lung A549 cells for a period of 24 hours. As seen in Fig.1 E, soot did not induce apoptosis, as soot exposure to cells did not cause any DNA fragmentation inside cell nucleus. Mechanistically, at molecular level to assess soot induced protection from apoptotic cell death, we measured cleaved caspase-3 expression (a marker of apoptosis). Fig. 1 F, shows that, soot exposure did not induce cleaved caspase-3 level in A549 cells in 24 hours of exposure period and minimal expression was comparable to control. This data further confirmed, previous results of DAPI staining (Fig. 1 E) that, soot does not induce apoptosis of A549 cells and may be involved in the cell proliferation. Subsequently, based on our speculation that, soot may work as a carbon source to these cells or inducing protective mechanisms, we have further tested effect of soot on the cells proliferation of A549 cells in glucose deprived (glucose free medium) conditions. Different concentrations (500 µg/ml, 1000 µg/ml, 2000 µg/ml) of soot were exposed to lung A549 cells for a period of 24 hours. As seen in Fig. 1 C, soot significantly induced cell proliferation of A549 cells as compared to control cells. Additionally, to assess autophagic cell death, we measured Light chain-3 (LC3) expression by seeing appearance of punctate staining in A549 cells in response to soot. As seen in the Fig. 1 G., soot exposure to cells did not cause LC3 expression in cells and thus confirmed that, soot did not cause any autophagic cell death in the A459, lung epithelial cells. Fullerene soot (2000 µg/ml) also induced cell proliferation in phosphate buffered saline alone (Sup Fig. 1).

**Fig. 1.**
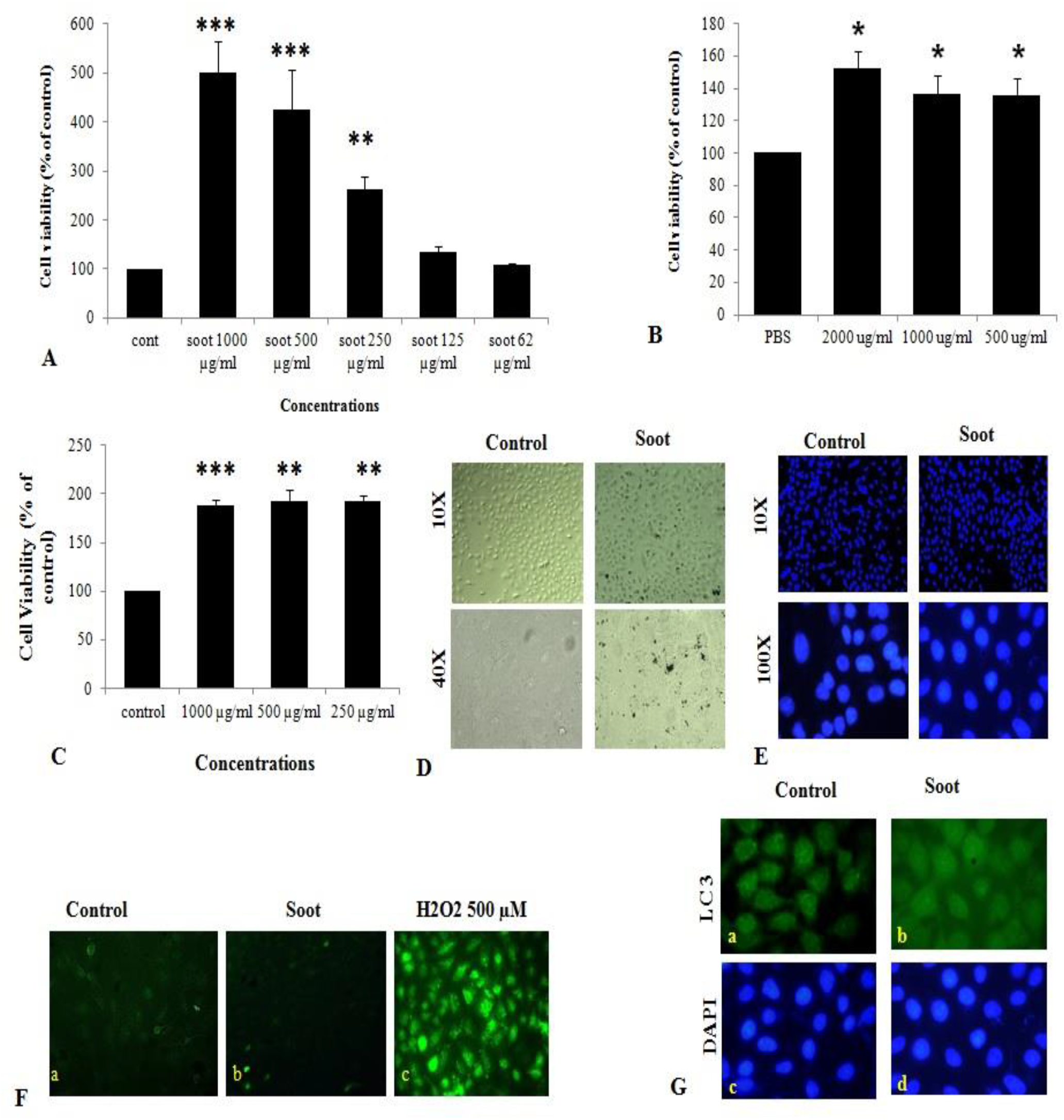
Effect of soot on cell proliferation of A549 cells. Histograms represent Mean ± SEM of different treatment groups. (A) Dose response of fullerene soot on the A549, cells in regular medium (DMEM:Ham’s F12+ 10% FBS) measured by MTT. (B) Dose response of fullerene soot (dispersed in the phosphate buffer saline), on proliferation of A549 cells was measured by MTT assay. Dose dependent study of fullerene soot on A549 cells in glucose free (GF) medium measured by kit (fluorescence method). (D) Bright field images of soot exposed A549, human lung epithelial cells. Top panel shows the lung epithelial A549 cells at 10x original magnification while lower panel shows the control and soot exposed cells at 40x original magnification. (E) DAPI stained control and soot exposed cells. The upper panel shows, A549 cells at 10x original magnification while lower panel shows control and soot exposed cells at 100x original magnification. (F) Cleaved caspases-3 expression in A549 cells in response to soot. Immunofluorescence photomicrographs of A549 cells are shown after 24 hours of H_2_O_2_ exposed cells. Green fluorescence shows the cleaved caspases-3 expression in A549 cells (at 40x original magnification). H_2_O_2_ 500 µM was used as a positive control for measurement of cleaved caspase-3. (G) LC3 punctate staining, in A549, cells. Lower panel shows that DAPI staining for the nucleus. Upper panel shows LC3 immunostaining. The upper and lower panel shows photograph at 100x original magnification. Green fluorescence shows the LC3 punctate staining. * p<0.05 treated versus control, ** p<0.01 treated versus control, ***=p<0.01 treated versus control. Data shown here is one representative experiment out of three.

### Fullerene soot inhibits autophagic and apoptotic cell death of human lung epithelial A549 cells

Fullerene soot did not induce cell death (apoptotic or autophagic) and causes cell proliferation of A549 cells. This suggested us that, soot may be inhibiting autophagy and thus inducing cell proliferation. To test this hypothesis that, “soot inhibits autophagy” we used LPS (Lipopolysaccharide) to induce autophagy in A549 cells and measured inhibition of LPS-induced autophagy due to soot. To accomplish this, A549 cells were exposed to LPS alone and in combination with fullerene soot. Measurement of autophagy was done by observing LC3 punctate staining. As seen in Fig. 2A, LPS profoundly induced autophagy in A549 cells by showing LC3 punctate staining. Fullerene soot inhibited LPS-induced punctate staining in A549 cells (Sup Fig. 2). The protein expression profile of autophagic genes were also measured in response to LPS alone or in combination with soot. Fullerene soot changed the expression profile of LPS-induced autophagic genes i.e. beclin-1 and ATG-5. Apart from autophagic genes, we have also tested the proliferation responsive genes in presence and absence of soot and LPS. As seen in the Fig. 2B, soot up regulated the expression profile of Ki-67 (a marker of cell proliferation) gene and thus shows that, soot induced proliferation.

**Fig. 2.**
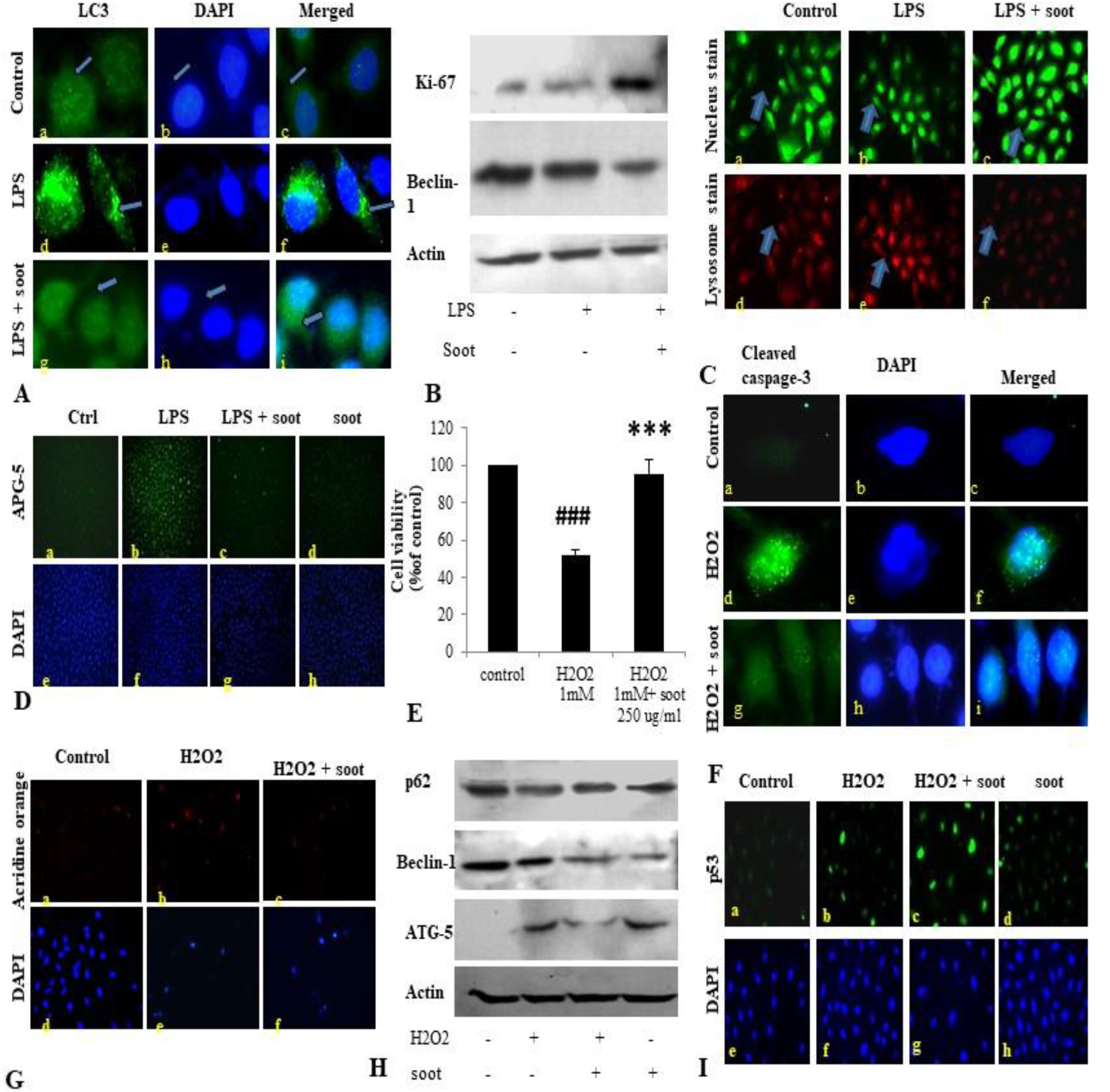
Fullerene soot inhibits LPS-induced autophagy and H_2_O_2_ induced cell death of A549, cells. (A) Effect of LPS on LC3 punctate staining in A549 cells in 24 hours exposure. a-c, immunofluorescence photomicrographs of control (no LPS) A549 cells are shown. Green fluorescence shows LC3 punctate staining (100x). d-f, LC3 punctate staining in LPS (10 µg/ml) treated cells. g-i, LC3 punctate staining in LPS + fullerene soot treated cells. (B) Western immunoblotting profile of soot on LPS-induced human lung epithelial cells. (C) Effect of soot on LPS induced acridine orange staining in A549 cells (at 40x). (D) Effect of fullerene soot on LPS induced ATG-5 expression profile in human lung epithelial cells (at 10x). (E) Effect of fullerene soot on the H_2_O_2_ induced cell death of A549 cells as measured by the MTT assay. (F) H_2_O_2_ induced cleaved caspage-3 expression is inhibited by the fullerene soot (100x). Left panel shows green fluorescence showing expression of cleaved caspage-3, middle row shows the counter staining by DAPI and right panel shows the merged images. (G) Acridine orange staining in response to H_2_O_2_ induced autophagic response (20x). H, Western immunoblotting profile of the proteins associated to the autophagic and apoptosis genes. (I), Expression profile of p53 in response to H_2_O_2_ and soot. Upper panel shows the p53 immunostaining and the lower panel shows the nuclear staining by the DAPI (20x).

In addition to autophagic cell death, we also measured inhibition of apoptosis by fullerene soot. To measure inhibition of apoptotic cell death we tested inhibition of H_2_O_2_ induced cell death by soot. To accomplish this, A549 cells were exposed to H_2_O_2_ (500 µM) alone and in combination with the soot (1000 µg/ml). Assessment of apoptosis was done by measuring expression profile of cleaved caspase-3, DAPI nuclear fragmentation staining and measurement of apoptotic associated proteins. As seen in Fig. 2 F, H_2_O_2_ significantly stimulated cleaved caspases-3 expression as measured by the immunocytochemistry and also showed nuclear fragmentation as seen by DAPI staining. Soot inhibited cleaved caspage-3 and nuclear fragmentation. The western immunoblotting profile shows that, soot inhibited expressions of autophagic and apoptosis genes stimulated by H_2_O_2_ exposure. As seen in the Fig. 2 H, soot changed the beclin-1 and ATG-5 gene expression showing H_2_O_2-_induced autophagy is inhibited by the fullerene soot exposure. This study confirmed that soot is a major inhibitor of autophagic and apoptotic cells death.

### Fullerene soot inhibits starvation induced autophagy and induces proliferation of A549 cells, which is reversed by Akt-Inhibitor X

It is well established that, glucose starvation of cells induces autophagy in cells (Nakai et al. 2020). Therefore, we tested whether soot treatment can inhibit starvation-induced autophagy. To accomplish this, A549 cells were starved of glucose for 24 hours and assessment of autophagy parameters were carried out in different conditions. As seen in the figure (Fig. 3), starvation of cells have significantly up regulated autophagy in lung epithelial (A549) cells as seen by the unregulated expression profile of autophagy genes beclin-1, p62, LC-3. Soot exposure has significantly inhibited the starvation-induced beclin-1 expression. Glucose starvation of the A549 cells has significantly up regulated p62 punctate staining in a 24 hours duration time period. Apart from autophagy genes, proliferation responsive genes ki-67 also up regulated due to soot exposure in a 24 hour period. Soot significantly inhibited autophagy in A549 cells as seen by the LC3 punctate staining.

**Fig. 3.**
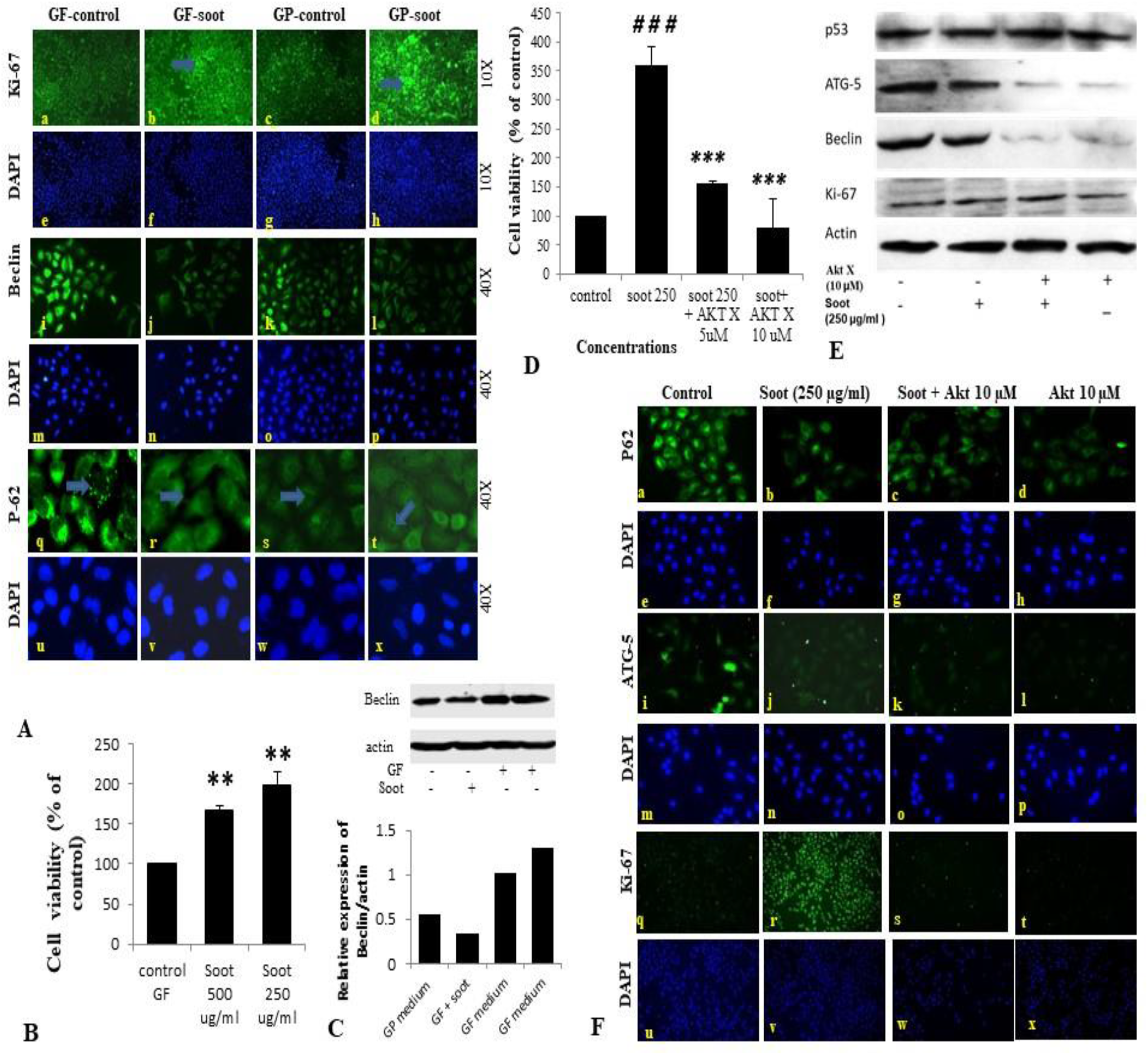
Fullerene-soot inhibited starvation induced autophagy in A549 cells, which is reversed by Akt inhibiter-X. (A) (a-d), expression profile of ki-67 (green fluorescence) in the presence and absence of soot in the mediums with and without glucose. a = glucose free (GF) medium without soot, b= glucose free medium with soot. c = medium with glucose. d= medium with-glucose with soot. (A) (i-l), expression profile of Beclin-1. i = glucose free medium, j= glucose free medium with soot. k= medium with glucose, l= medium with-glucose after soot exposure. (A) (q-t), p-62 localization profile after the exposure of fullerene soot in glucose free and glucose containing medium. q= glucose free medium, r= free medium with soot. s= medium with glucose, t= fullerene soot exposure in medium with-glucose. (B) Cell proliferation assay in the presence and absence of fullerene soot in glucose free medium. ** p<0.01 compared with control (C), western immunoblotting of beclin due to exposure of soot in the presence and absence of glucose (top), histograms representing densitometric intensity of bands (down). (D), Effect of Akt inhibitor X on the cell proliferation of A549 cells due to soot in glucose free medium. ### p<0.01 compared with control and *** p<0.001 compared with fullerene soot treated group. (E) Western immunoblotting of autophagic proteins in response to Akt inhibitor X. (F), Expression profiles of genes in glucose free medium, after treatment with Akt inhibitor X. Expression profile of p62 (20x). a= glucose free medium b= medium with soot. c= soot exposure with Akt inhibitor X (5 µM), d = soot with Akt inhibitor X (10 µM). ATG-5 expression profiles (20x), (i-l). i= glucose free (GF) medium j= medium with soot. k= soot exposure with Akt inhibitor (5 µM), l= soot exposure with Akt inhibitor X (10 µM). Ki-67 expression profile (10x). q= glucose free medium, r = medium with soot. S = fullerene soot exposure with Akt inhibitor X (5 µM), t= soot exposure with Akt inhibitor X (10 µM). DAPI staining (blue image) is shown blow respectively of each green fluorescence image.

Akt signalling pathway is associated with the starvation-induced autophagic response but the role of soot on Akt pathway activation was not known. To understand the mechanism of soot associated cell proliferation and autophagic inhibition, we have tested the role of Akt inhibitor X on soot-induced cell proliferation and autophagic response. Cells were treated with different concentrations of soot in the absence of glucose and effect of Akt inhibitor X was measured on the cell proliferation and autophagic response. Soot exposure significantly stimulated cell proliferation of human lung epithelial cells A549 in 24 hours. As demonstrated in Fig. 3 D, it was seen that Akt inhibitor X (5 µM and 10 µM) has significantly inhibited soot induced cell proliferation of human lung epithelial cells in a dose dependent manner. In addition to the inhibition of cell proliferation, Akt inhibitor also showed down regulation of expressions of autophagy genes i.e. p62, beclin-1 and ATG-5 in a dose dependent manner. Soot exposure also inhibited p62 punctate staining in the cells showing inhibition of autophagic flux.

Further to understand the effect of ambient soot, effect of diesel soot (diesel exhaust particles or DEP) and biomass burnt (BI) soot (wood burning) was evaluated on A549 cells. As seen in Fig. 4 A, biomass burning soot did not cause cell death of lung airway epithelial cells A549. Soot concentration (250, 500, and 1000 µg/ml) increased MTT score at 24 hours showing cell proliferation of lung epithelial cells A549. Similarly, we also tested effect of diesel exhaust particles. Lung A549 cells were exposed to different concentrations (250 µg/ml, 500, µg/ml and 1000 µg/ml) of DEP for a period of 24 hours. As seen in Fig. 4 B, DEP significantly caused cell proliferation of A549 cells as compared with control cells. In addition to cell proliferation, we also tested expression profiles of autophagic genes in response to DEP and biomass burnt soot. As shown in Fig. 4 F, soot exposure to A549 cells significantly up-regulated the expressions of autophagic genes in response to both types of soot. We have also tested the role of Akt inhibitor X on DEP and BI induced cell proliferation and autophagic responses. Cells were treated with different concentration of DEP,BI and in the absence of glucose and effect of Akt inhibitor X was measured on the cell proliferation and autophagic responses. DEP and BI soots exposure to the A549 cells significantly stimulated cell proliferationin 24h hr. Interestingly, as demonstrated in Fig. 4C, Akt inhibitor X (10 µM) did not inhibit the DEP induced cell proliferation of human lung epithelial cells. In fact, Akt inhibitor treatment significantly potentiated the DEP-induced cell proliferation. Co-exposure of Akt-Inhibitor X with DEP also showed increase in the expression of autophagy gene beclin-1 in a dose dependent manner. We got similar results with the COX-2 inhibiter Etoricoxib, to that of Akt-inhibiter X. Etoricoxib also did not down-regulated DEP-induced cell proliferation of A549 cells. Inhibition of COX-2 affects endoplasmic reticulum mediated autophagy. However, chloroquine, an inhibiter of autophagic flux, has significantly inhibited DEP induced cell proliferation of A549 cells (Fig. 4). Similarly, autophagic inhibitor 3-MA has also significantly inhibited DEP induced cell proliferation in dose dependent manner (Sup Fig. 4).

**Fig. 4.**
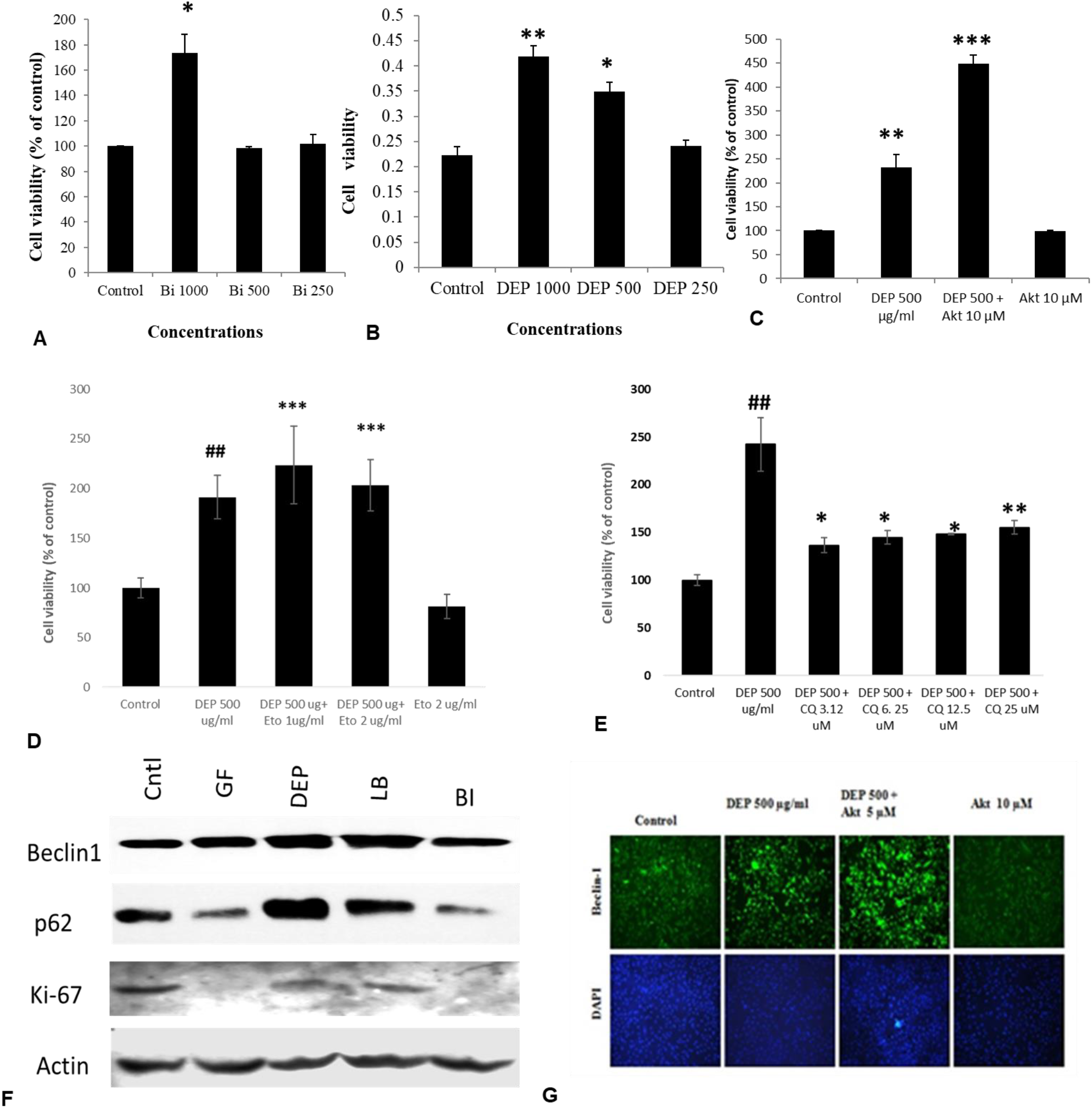
Dose dependent study of DEP (diesel exhaust particles) and biomass burning soot on the cell proliferation of A549 cells; effect of COX-2 inhibiter, autophagic inhibiter chloroquine and Akt Inhibitor X. Histograms represent the Mean ± SEM of the different treatment groups. Dose response of biomass burning soot on the A549 human lung epithelial cells in glucose free medium measured by MTT assay. (B) Dose response of diesel exhausts particles, on A549 human lung epithelial cells in glucose free medium measured by MTT assay. (C) Effect of Akt inhibitor X, on the diesel exhaust particle induced cell proliferation. (D). Effect of Etoricoxib (a COX-2, inhibiter) on the DEP induced cell proliferation of A549 cells. (E). Effect of chloroquine (an inhibiter of autophagic flux) on the DEP induced cell proliferation of A549 cells. (F) Western immunoblotting profile of different autophagic proteins in response to different kinds of soots, i.e., diesel exhaust particles, soot from biomass burning. (E) Effect of Akt inhibitor X, on the diesel exhaust particle induced beclin-1 expression profile.* p<0.05, ** p<0.01 and *** p<0.001 compared with control in figures A, B, C. # p<0.05, # p<0.01 compared with control, and * p<0.05, ** p<0.01 and *** p<0.001 compared with DEP or soot treated group in figures D and E.

### Fullerene soot particles are internalized and co-localizes with autophagosomes associated protein p62

We speculated that, soot associated autophagic response is mediated by its physical presence with the autophagosomes. To test this speculation, we have first coated 6 well plates with methanol treated fullerene soot and allowed the cell to grow on the coated plates. After the incubation period of 24 hours, the cells were fixed and seen in the fluorescence microscope. As seen in Fig. 5 B., cell has internalized some particles of fullerene soot (reflected in the form of red fluorescence). To measure the interaction of these fullerene soot particles we also tested p62 expression and localization pattern in the soot treated cells. Fig. 5 B shows that, p62 co-localizes with the internalized soot particles. To assure that treatment with the methanol did not alter the cell proliferation associated effect of the lung epithelial cells we have also tested the effect of methanol treated soot on the cell proliferation of A549 cells. The methanol treated soot concentration 250 µg/ml, 500 µg/ml, and 1000 µg/ml has increased the MTT score at 24 hours of incubation period. The data in the Fig. 5. denotes that, soot in fact caused the cell proliferation of lung epithelial A549 cells. It was interesting to note that, soot particles co-localise with the autophagy proteins suggesting its role in autophagy.

**Fig. 5.**
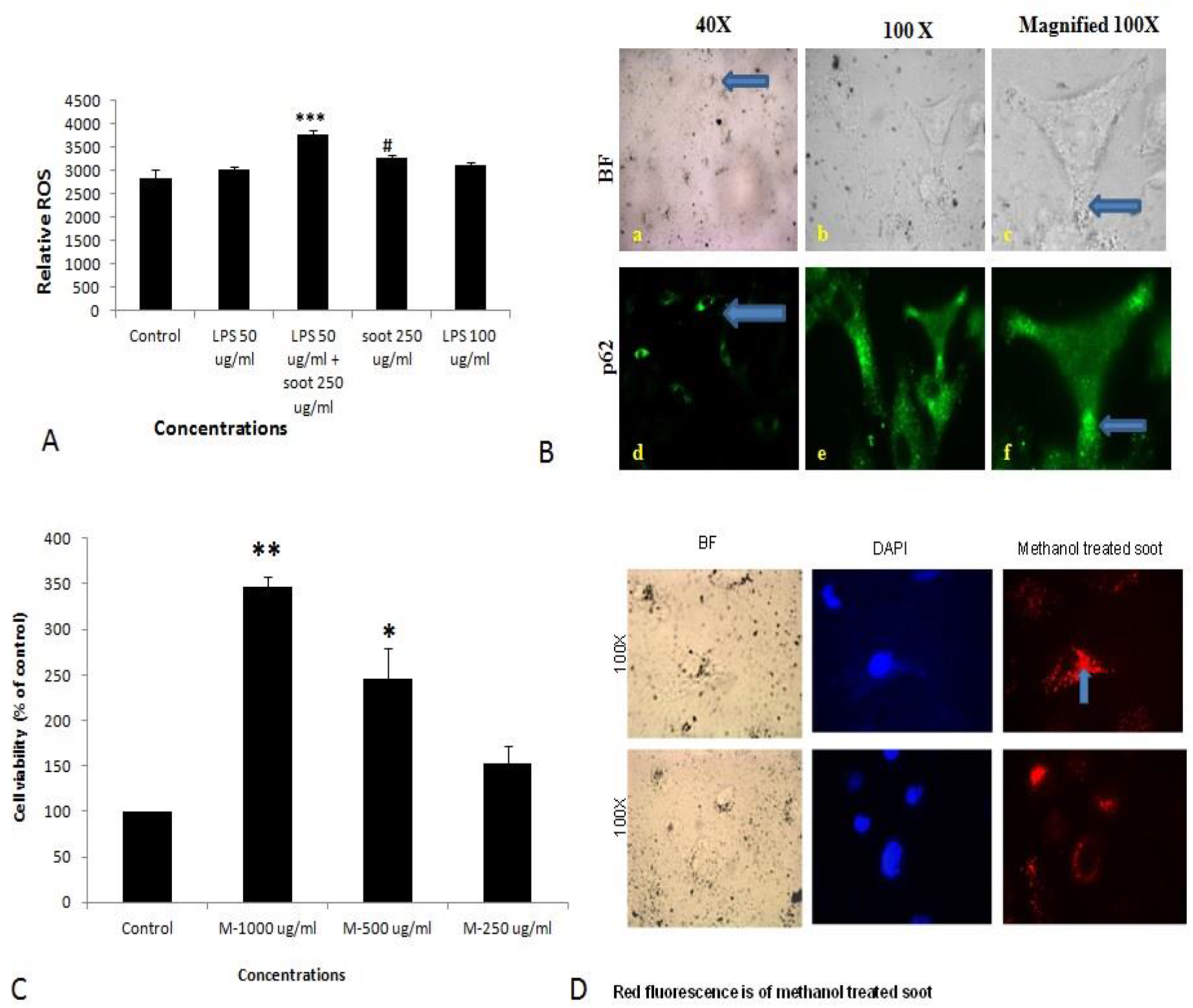
Fullerene soot enters the cell and co-localize with the autophagic protein p62. (A) Histograms represent comparative ROS formation in the A549 cells. *** p<0.001 compared with control and # p<0.05 compared with LPS + fullerene treated group. (B) Cellular localization of soot with the autophagy protein p62. Green fluorescence images show the p62 localization. Photomicrographs were captured at 40x, and 100x original magnifications. The upper panel corresponds to the bright field images and the lower panel shows the p62 localization as a green fluorescence. (C) Dose dependent study of the methanol treated fullerene soot on the cell viability of A549 cells. .* p<0.05, ** p<0.01 compared with control. (D) Mmeasurements of internalization pattern of soot inside the cell. Red fluorescence shows the methanol treated soot inside the cell. The corresponding DAPI photographs are also shown with the photograph to position the cell nucleolus.

### Soot induces cell proliferation and inhibits autophagy in the in vivo model of mice

To further clarify that, soot inhibited autophagy in the *in vivo* conditions, we tested effect of fullerene soot on autophagic response when it is intranasally delivered in mice. To accomplish this, 100 µg of soot was intranasally delivered to mice. A total of two intranasal challenges were given to mice in 12 days with an interval of 6 days. (exposure plan of mice is depicted in fig. 6 A) Control mice were delivered with the saline alone. As seen in the Fig. 6 C, basal autophagic markers were inhibited by the soot challenge. As seen in the Fig. 6 C, fullerene soot has significantly upregulated ki-67 expression in lung airway epithelium giving confirmation of the proliferation inducing property of fullerene soot. Fullerene soot also down regulated autophagy responsive genes beclin-1 in lungs and in the oesophagus, depicting its important autophagy inhibiting properties. In addition to beclin-1 gene expression, fullerene soot has also down regulated the expressions of autophagy proteins LC3, p62 and APG-5 in the lung (Fig. 6 D). These results suggest that, fullerene soot inhibits autophagy and induces cell proliferation also in *in vivo* conditions in healthy mice.

**Fig. 6.**
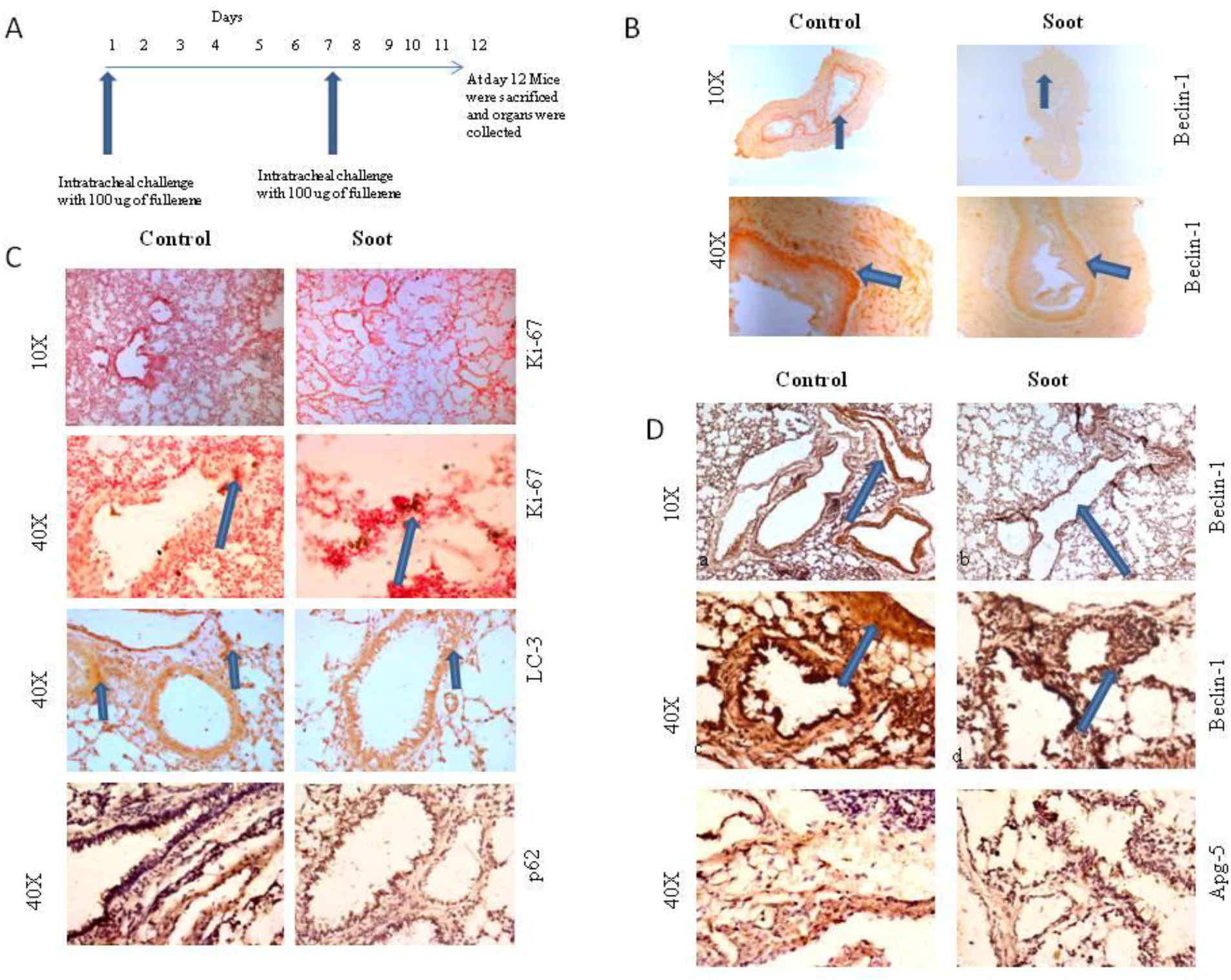
Assessment of soot induced autophagic response in the mice. (A) The schematic representation of challenge to the mice with fullerene soot. Vertical arrows show the day of exposure to the mice. The horizontal line shows number of days of progression of experiment. beclin-1expression in the esophagus of mice. a-, esophagus section of saline treated mice b, esophagus section of fullerene soot treated mice. (C) Ki-67 and LC3 expression pattern in the lung’s section of saline treated and in the soot treated mice. (D) beclin-1 expression in the lung’s section of mice. a, section of saline treated mice b, esophagus section of fullerene soot treated mice, c and d are the same at 40x magnification. ATG-5 gene expression profile in the lung and in the esophagus of saline and soot treated mice.

### Effect of fullerene nanoparticles (agglomerates) on the exprerssion profile of autophagy genes (in HEK 293), organic carbon content and p53 traslocation

A549 is the one of the immortalised cancerous cell lines and does not represent the characteristics of a normal cell type. Therefore, to understand the effect of fullere soot, HEK293 (non cancerus cell line) was used as a non cancerous cell line. Furthermore, the effect of fullerene soot on expressions of autophagic genes in HEK293 cells (Fig. 7) was also assesed. As seen in the Fig. 7 A and B, fullerene soot successfully attenuated autophagic genes beclin and ATG-5 expressions (in glucose free medium). It also illustrates that, fullerene soot exposure did not cause any traslocation of p53 protein in HEK293 cells depicting its non-carcinogenic effect (Fig. 7C). In addition to this, the cell proliferation effect of fullerene soot on the SH-SY5Y cells was also tested, which showed, fullerene soot significantly induced cell proliferation of human neuroblastoma, SH-SY5Y cells. Different concentration of the soots did not cause cell death of human SH-SY5Y cells. As described in the previos results, fullerene soot induces cell proliferation and supreses autophagy, it was desired to know that, whether fullerene soot particles are being used as a carbon source directly or indirectly by cells. To test this, organic carbon content (OC) at elemental carbon (EC) level was measured using ECOC instrument. As shown in Fig. 7E, organic carbon content of fullerene soot treated cells increased with time as measured at different time intervals Fig. 7E. This may indicate that elemental carbon of fullerene soot particles was utilised by cells as a carbon source. This may explains the mechanism of survival and cell proliferation of different cell types due to fullerene soot. The data on the disagregation of fullerene aggluromarate also support that soot particles are dissociated at a higher rate with time in medium (Fig. 7F).

**Fig. 7.**
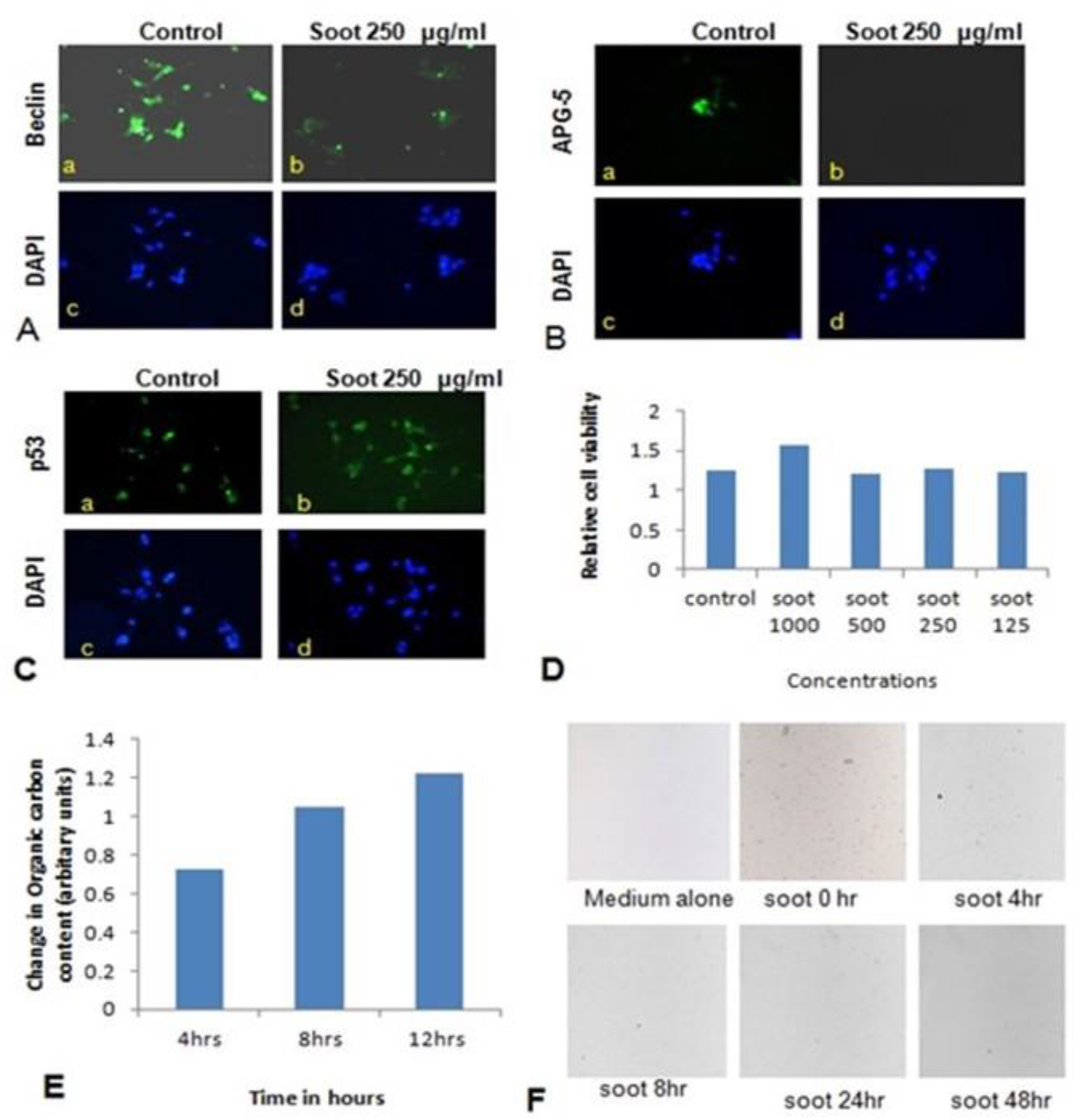
Effect of fullerene nanoparticles (agglomerates) on the expressions profile of autophagy genes (in HEK-293), organic carbon content and p53 translocation. A and B; effect of fullerene soot in HEK-293 cells on beclin and ATG-5 expressions (in glucose free medium). C; effect of fullerene soot on traslocation of p53 protein in HEK-293 cells. D; Effects of different concentartions of fullerene soot on cell proliferation of human neuroblastoma cells, SH-SY5Y cells. E; organic carbon content at elemental level using ECOC instrument was analysed. F; The data on the disagregation of fullerene aggluromarates with time.

## Discussion

Exposure of environmental soots were thought to cause severe lung pathologies, however their underlying mechanisms were not properly understood (Niranjan, Thakur 2017). In the present study, we demonstrate effects of fullerene soot and DEP (diesel exhaust partilcles) on the proliferatiion of human lung epithelial and other cells. We show that, both kinds of soot induce proliferation of lung epithelial cells via interruption of distinct autophagic mechanisms (Pidsley et al. 2018). Interestingly, soots did not induce cell death of lung epithelial, A549 cells. In fact, soots not only induced proliferation of lung epithelial A549 cells in *in vitro* culture conditions but also in mice (Shin et al. 2013). At the molecular levels, both soots regulate different autophagic mechanisms to induce proliferation of lung epithelial cells. Our work, to dissect out distinct mechanisms of autophagy to regulate cell proliferation in response to exposure of soots provide an important insight in the pathological mechanisms of many soot-induced disorders.

It is important to know that, autophagy is involved in the regulation of cell proliferation of many cell types. Autophagy regulator beclin-1, critically controls cell proliferation during tumerigenesis (is it important to bring tomerogensis, what about normal cell division) (Cicchini et al. 2014). Apatinib, a tyrosine inhibitor decreases cell proliferation and initiate autophagy via the PI3K/Akt/mTOR pathway, establishing a link in the cell proliferation and autophagy regulation (Meng et al. 2020). Similarly, autophagy dependent proliferation was seen in response to microRNAs-107 in breast cancer cells (Ai et al. 2018) (putting cancer in the start of discussion may not be right). The present findings also align with these existing studies (Fig.1 and Fig. 6) with multiple evidences both at cellular and molecular levels. In this study, autophagy is inhibited by fullerene soot which may be a mechanism behind the induction of proliferation of lung epithelial cells leading to cell hyperplasia thus resulting in lung dysfunctions (Weng et al. 2014; Qiang et al. 2014). Autophagy and cell proliferation is also linked with the morphological changes and apoptosis of cells which we also found in response to soots (Chen et al. 2020). At molecular level, soot did not induce cleaved caspase-3 expression (Fig. 1, F), confirming its inability to induce type-I cell death (apoptosis) of the human lung epithelial cells (Atale et al. 2014; Han et al. 2014). We also observed that, soot did not cause DNA fragmentation in human lung epithelial cells and thus did not induce cell death (Gai et al. 2014).

One of the important conclusions that emanates from current work is the fact that, both soots do not cause cell death, which is evidenced by many lines of experiments (Fig 1 and (Fig. 2). In fact it indicated their protective effects aginsts cell death inducers. LPS (Lipopolysaccharide) is an endotoxin and known to induce autophagy in different cells including lung epithelial cells by toll like receptors (Pei et al. 2015; Waltz et al. 2011). Recently, it was shown that inhibition of LPS induced autophagic response is linked with the cell proliferation in macrophage (Jakhar et al. 2014). Similarly, in the present study, LPS has significantly up-regulated LC3 punctate staining in the cells showing induction of autophagy. Treatment with fullerene soot clearly inhibited LPS-induced autophagic response and enhanced cell proliferation, showing the role of autophagy in proliferation. In addition to this, fullerene soot has significantly up regulated the Ki-67 expression profile confirming its proliferative potential at molecular level (Troncoso et al. 2012). Hydrogen peroxide is known to cause apoptosis in many cells. H_2_O_2_ exerts its effect mainly by producing oxidative stress response. The oxidative stress may or may not be associated with the autophagic response of cells (Zhang et al. 2015; Chen et al. 2008). In the present study, H_2_O_2_ has significantly induced cell death by the way of autophagy and apoptotic mechanism in human lung epithelial cells. Fullerene soot significantly inhibited H_2_O_2_ induced apoptosis and autophagic death. As shown in the Fig. 2 D, fullerene soot also reversed H_2_O_2_ induced autophagy proteins beclin-1, p62, ATG-5 showing its important mechanism of action but not by its antioxidant property As seen in the result section (Fig. 5 A) that soot does not show its antioxidant properties at the used concentrations. It is clearly seen that, H_2_O_2_ completed the autophagy and initiated the apoptotic events however soot alone did not induce autophagy. It is already known that, H_2_O_2_ induces apoptosis via way of cleaved caspase-3 induction (Li et al. 2015; Back et al. 2015). In this study, soot has inhibited H_2_O_2_ induced apoptosis and also inhibited cleaved caspase-3 expression and nuclear localization suggesting its role at transcription factor level. (Antioxidatnt lines needs refinements, some repetetions are seen)

Autophagic cell death is an important mechanism in nanoparticles toxicity (Loos et al. 2014; Chen et al. 2014). NLRP3 inflammasome and autophagy activation by metal nanoparticles was found to suppress cell proliferation (Sasabe et al. 2020). These metal nanoparticles (gold, silver, and palladium) also caused lysosomal dysfunctions suggesting an important role of autophagy bynanomaterials. Autophagic protein LC3 is associated with toll like receptor signalling pathway of cells (Acharya et al. 2016). These studies indicated that, soots being nanonaprticles may be affecting autophagy mechanism and thus inducing cell proliferation. It is noted that, cellular accumulation of silica nanoparticles cause lysosomal dysfunctions and colocalize with the autophagic proteins (Schutz et al. 2016). Surface properties and core stability of nanoparticles were found to be responsible for colocalization with autophagyc molecules and trigger mTOR pathway in hepatocellular cells (Lunova et al. 2017). We observed that, fullerene soot particle are internalised and colocalise with the autophagic protein p62 which refelects an important role of autophagy in the trigerring of soot particle degradation or digestion (Fig. 5B). Considering the nanoparticle mediated effects of soot, we further tested autophagic and apoptosis inhibitory properties of fullerene soot for mechanistic understanding.

To understand the autophagic mechanism of soot mediated regulation of cell proliferation, we starved A549 cells and triggered starvation-induced autophagy (Mofarrahi et al. 2013; Huang et al. 2013). PI3/Akt signalling pathway is a key regulatory mechanism involved in autophagic response induced by starvation (Wang et al. 2013; Wu et al. 2009). Akt induction is a survival pathway and inhibition of Akt is linked with the inhibition of cell proliferation and induction of cell death (Bokobza et al. 2014). As depicted in the results, starvation of cells to glucose induces autophagy in A549 cells which was significantly inhibited by the treatment of fullerene soot. This indicated that, fullerene soot might be utilised as an energy source and compensated the energy associated demand of the starved cells. As shown in the Fig. 3 E, fullerene soot-induced cell proliferation is significantly inhibited by the Akt inhibitor X, suggesting that, it is an Akt dependent pathway. It is important to note that, apart from cell proliferation, Akt Inhibitor X also decreased the expressions of autophagy genes showing the strong evidence that fullerene soot inhibits induction of autophagy at very initial points (Fig. 8). Furthermore, it is evidenced that activation of Akt mediated autophagy inhibits apoptosis induced by DRAM-mediated mitophagy (Liu et al. 2014). Therefore, it is believed that fullerene soot might have inhibited apoptosis of A549 cells by a similar mechanisms. We have also shown that, fullerene soot is converted into organic carbon as measured by the OCEC instrument. Here for the first time, we report that, elemental carbon is converted into organic carbon however the detailed mechanism is still a subject of further investigation (Fig.7).

**Fig. 8.**
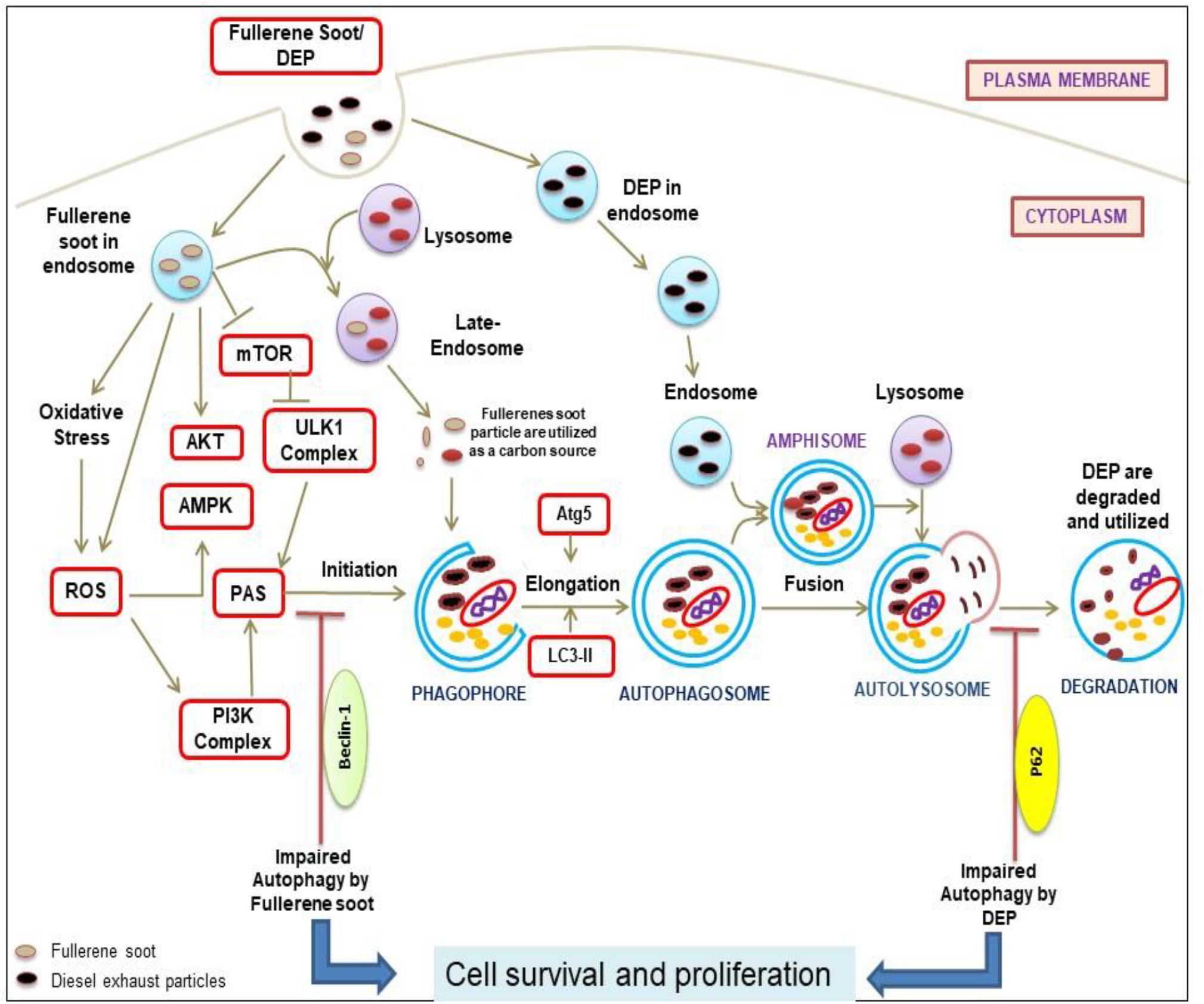
Diagramatic repregentation of proposed mechanisms of action due to soots (fullerene soot and DEP) mediated inhibitions of autophagy and subsequent induction of cell proliferation. As seen in the figure fullere soot inhibit autophagy at the level of beclin as they are directly utilized as an energy source. However, DEP inhibit the autophagic flux and does not allow the autophagy to complete possibly by regulating the expression profile of p62.

Similar to the fullerene soot, diesel exhaust particles (DEP) also induce cell proliferation of lung epithelial cells but did not down-regulate autophagic response (Gai et al. 2014). In fact, diesel exhausts particles increased autophagic response in human lung epithelial cells. Recently, it was shown that autophagy inhibition by Akt inhibitor is liked with inhibition of cell proliferation (Zhao et al. 2016). Impairment of autophagic flux was connected with the cell death, however the autophagic flux associated effect on cell proliferation of lung epithelial cells is still less clearly known (Yun et al. 2014; Masschelein et al. 2014). Here we support the fact that lysosomes trap lots of fullerene soot particles making it unable to receive cargo from other autophagosomes. This may be due to defective fusion of these organelles with lysosomes. As evidenced by in the results (Fig. 4 F) that, non-degraded p62 refelect the impaird autophagic flux in DEP treated cells. Based on the results we can only speculate that lysosomes maybe fusion-defective and thus inhibiting autophagy in DEP treated cells.

Existing literature suggests that, there are Akt dependent and Akt independent pathways (Kulik, Weber 1998; Marsh Durban et al. 2013). DEP inhibited autophagic flux and this might lead to the cell proliferation of airway epithelial cells (Riahi et al. 2016). Interestingly, it was observed that Akt inhibitor did not down-regulate the diesel exhaust particle induced cell proliferation. Indeed, Akt inhibitor X further potentiated the DEP-induced cell proliferation. DEP-induced autophagic response was also potentiated by the Akt inhibitor X. Here, it became clear that, dfferent autophagic mechanisms of cell proliferation exists in response to DEP (p62 expression up) and in fullerene soot (p62 expression down). These findings clearly show that DEP-induced cell proliferation and autophagic response is entirely different and does not depends on Akt pathway (Memmott, Dennis 2009). A previous study described that, BRAFV600E a molecule coordinates to the PI3K signaling to control melanoma cell proliferation which is independent of AKT signaling and supported the existance of Akt independent pathway in the regulation of cell proliferation (Silva et al. 2014). This study also strongly supports our findings indicating that, DEP may be following Akt independent pathways for the regulation of cell proliferation. Notably, DEP induced cell proliferation is inhibited by autophagic inhibiter 3-MA and chloroquine but not by COX-2 inhibiter “Etoricoxib” again pointed towards different mechanism of autophagy (Kaewjumpol et al. 2018). Chloroquine and 3-MA show inhibition of cell proliferation in a number of cell types providing a support to the present findings (Jiang et al. 2008; Fan et al. 2006; Dong et al. 2019).

In conclusion, it can be said that, autophagy is a key regulatory mechanism responsible for the cell proliferation in response to exposure of soots. The present finding of autophagic mechansism for the control of cell proliferation may throw light on the pathophysiology of tissue remodelling of severe diseases and on prospects of regenerative capabilities of fullerene soot. As the problems of respiratory dysfunctions due to air pollution are increasing throughout the world, proper knowledge about the pathophysiological processes would be a tremendrous advantage. Fullerene soot down-regulated autophagy and thus is able to cause cell proliferation of lung and other cell types. This cell proliferation (hyperplasia) may initiate the development of tissue remodelling affecting asthmatic pathology. In this study, the effect of fullerene soot is because of its chemical composition and structural properties rather than its antioxidant effects. Similar to the fullerene soot, the DEP also caused cell proliferation of cells but did not down regulate autophagic response. These results suggest that, more than one type of autophagic mechanisms operate in response to different types of soots. We also highlight the importance of fullerene soot, as a regenerative medicine to be developed and which can be used in the management of autophagy associated disorders. Irrespective of encouraging results which may be of clinical relevance a reassessment of study may be required to further strengthen the findings.

## Acknowledgements

AKT acknowledges the MHRD, Government of India for funding (MHRD/CESE/2013357) this interdisciplinary project to Prof. Sachchida Nand Tripathi, Prof. Tarun Gupta, Prof. Ketan Rajawat, Prof. Debajyoti Paul, and Prof. Ashwani Thakur. This study is part of this interdisciplinary project. The “young scientist grant No. SB/YS/LS-198/2014 to Dr. Rituraj Niranjan by SERB (DST, India.) is gratefully acknowledged.

## Author Contributions

*<u>Dr. Rituraj Niranjan</u>*, performed all experiments in cell culture and in mice model. He also did analysis and interpretation of data and wrote the manuscript. *<u>Kaushal Prasad Mishra</u>*, provided his assistance during the experimentations. *<u>Ashwani Kumar Thakur</u>* contributed in concept and design, analysis and critically reviewing and correcting of the manuscript, study supervision, and funding support.

## Abbreviations

DEP: Diesel exhaust particles;
COX-2: Cyclooxygenase-2;
DCF-DA: dichloro-dihydro-fluorescein diacetate;
LPS: Lipo-polysaccharide;
Akt: Protein kinase B;
LC3: Light chain 3;
ATG-5: Autophagy protein-5;
p62/SQSTM1: Sequestosome 1;
AMPK: Adenosine monophosphate-activated protein kinase;
GF: Glucose free;
GP: Glucose plus;
AD: aerodynamic diameter;
PAHs: polycyclic aromatic hydrocarbons;
ROS: Reactive oxygen species;
HRP: Horse Radis Peroxidase;
PM: particulate matter;
OCEC: Organic Carbon/Elemental Carbon Analyser;
HBSS: Hank’s balanced salt solution;
3-MA: 3-Methyladenine;
DMEM: Dulbecco′s Modified Eagle′s Medium.

## Supplementary information

### Results

#### 1. Fullerene soot induced cell proliferation of A549 cells in phosphate buffered saline (PBS)

Fullerene soot increased cell proliferation of A549 cells in absence of regular DMEM medium. In order, to assess the effect of fullerene soot as a carbon source and as cell proliferating agents we have tested its effect on cell proliferation of human lung epithelial cell line at higher concentration (2000 µg/ml). As seen in the figure, fullerene soot 2000 µg/ml has significantly increased the cell proliferation of human cells in 48 hours.

**Supplementary figure 1:**
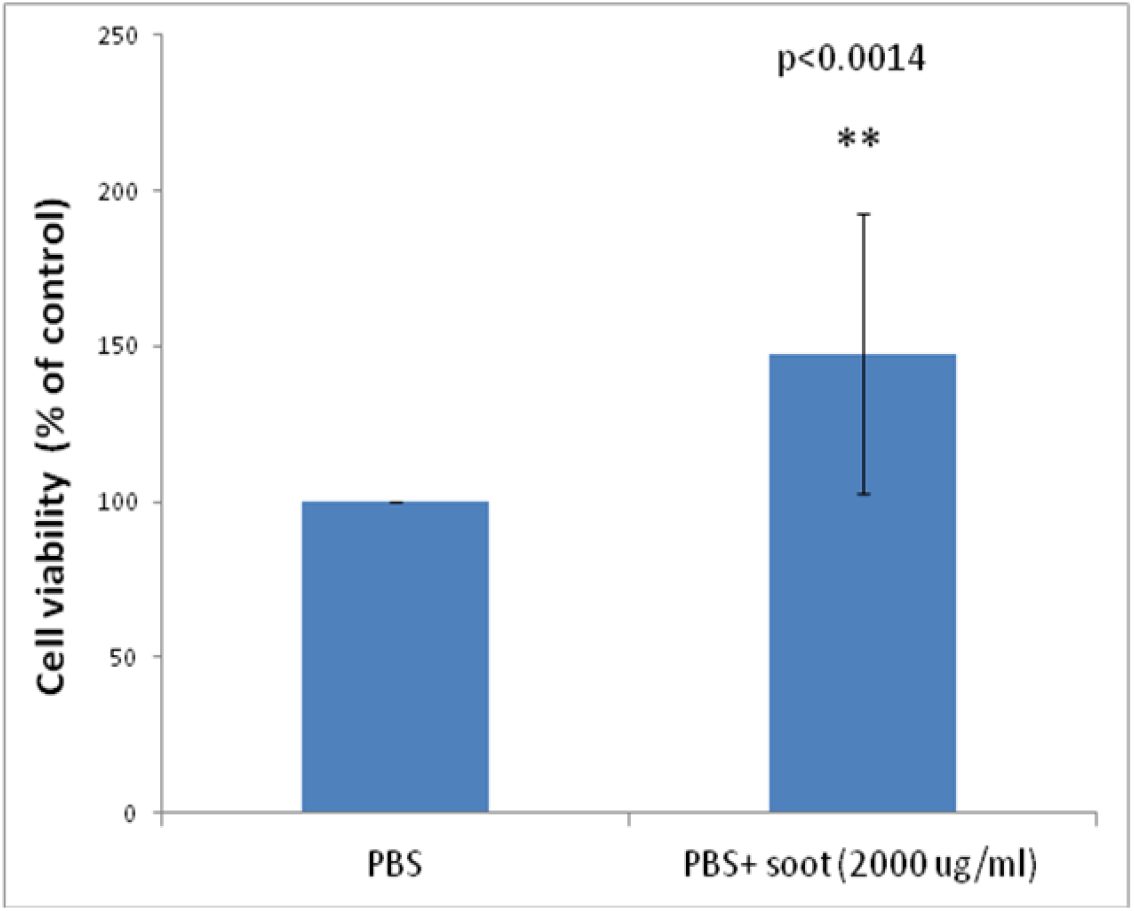
Cell proliferation of A549 cells in PBS. Histograms represent the Mean ± SEM of the different treatment groups. **=p<0.001 treated versus control.

#### 2. Fullerene soot inhibits LPS induced LC3 expression in A549 cells

It is known that LC3 is the marker for the autophagy. We checked the expression profile of LC3 in response to soot. As shown in the figure LPS has significantly up regulated expression profile of LC3 as seen by the punctate staining in the cells (middle panel). Exposure of fullerene soot has significantly inhibited LPS induced induction of LC3 expression in a 24 hour of exposure period.

**Supplementary figure 2:**
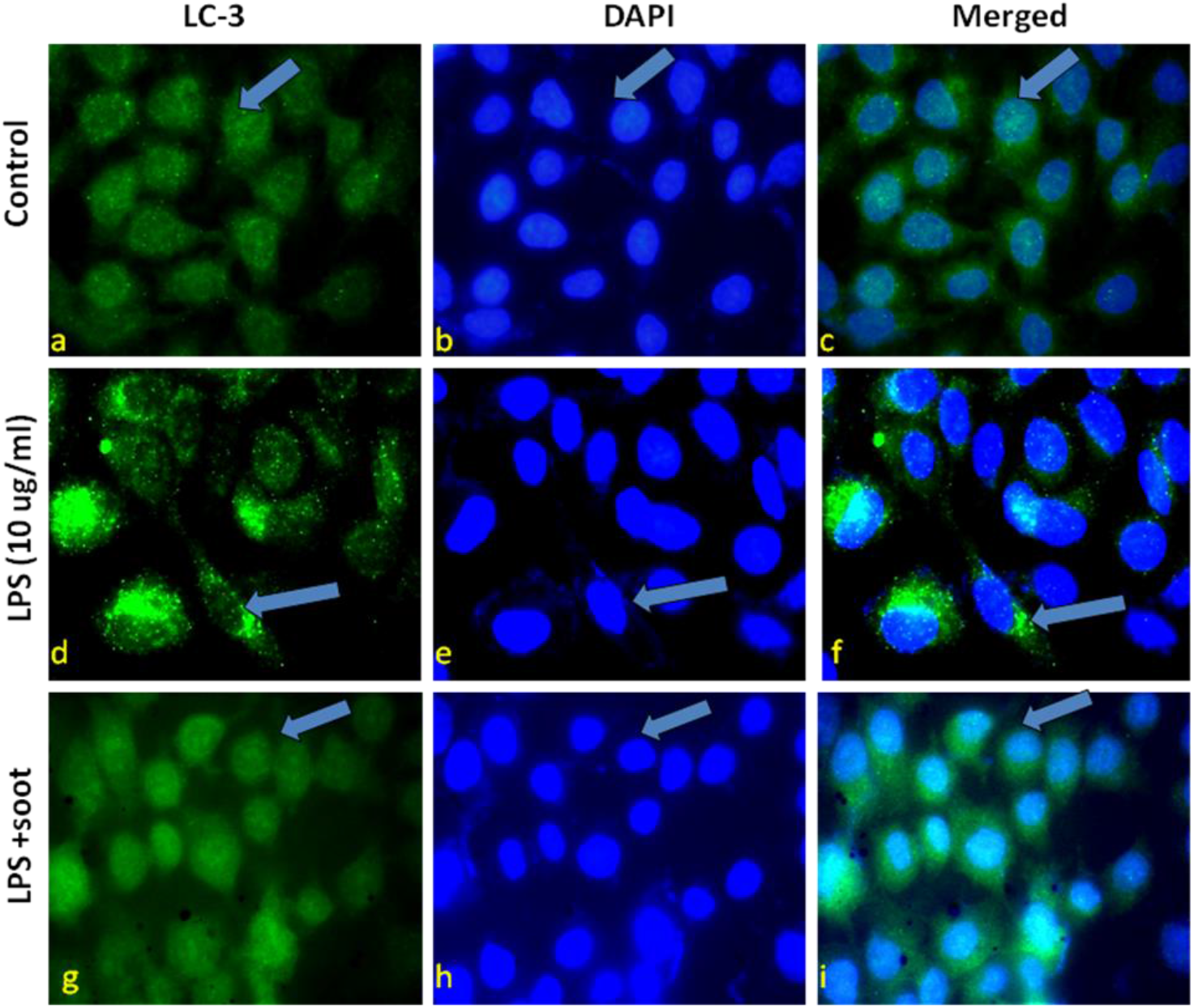
Soot inhibits the LPS induced autophagy in lung epithelial cells. A. LC3 punctate staining in control A549 cells. a-c, immunofluorescence photomicrographs of A549 cells are shown after the 24hrs of control (no LPS) cells. Green fluorescence shows the LC3 punctate staining. panel shows photograph at 100X original modification. d-f, LC3 punctate staining in LPS treated A549 cells. Immunofluorescence photomicrographs of A549 cells are shown after the 24hrs of LPS (10 µg/ml) exposed cells. g-i, LC3 punctate staining in LPS + soot (250 µg/ml) treated A549 cells.

#### 3. The profile of total protein content in response to fullerene soot and H_2_O_2_

The expression profile total protein content in response to H_2_O_2_ and soot in A549 cells was measured by the BCA protein estimation kit. To total protein expression is expressed in µg/ml. As shown in the figure that, H_2_O_2_ has increased the total protein content by inducing a set of cells signaling pathways associated to cell death. Fullerene soot decreased the H_2_O_2_ induced cell signaling pathways in dose depending manner indicating its antioxidative stress properties.

**Supplementary figure 3:**
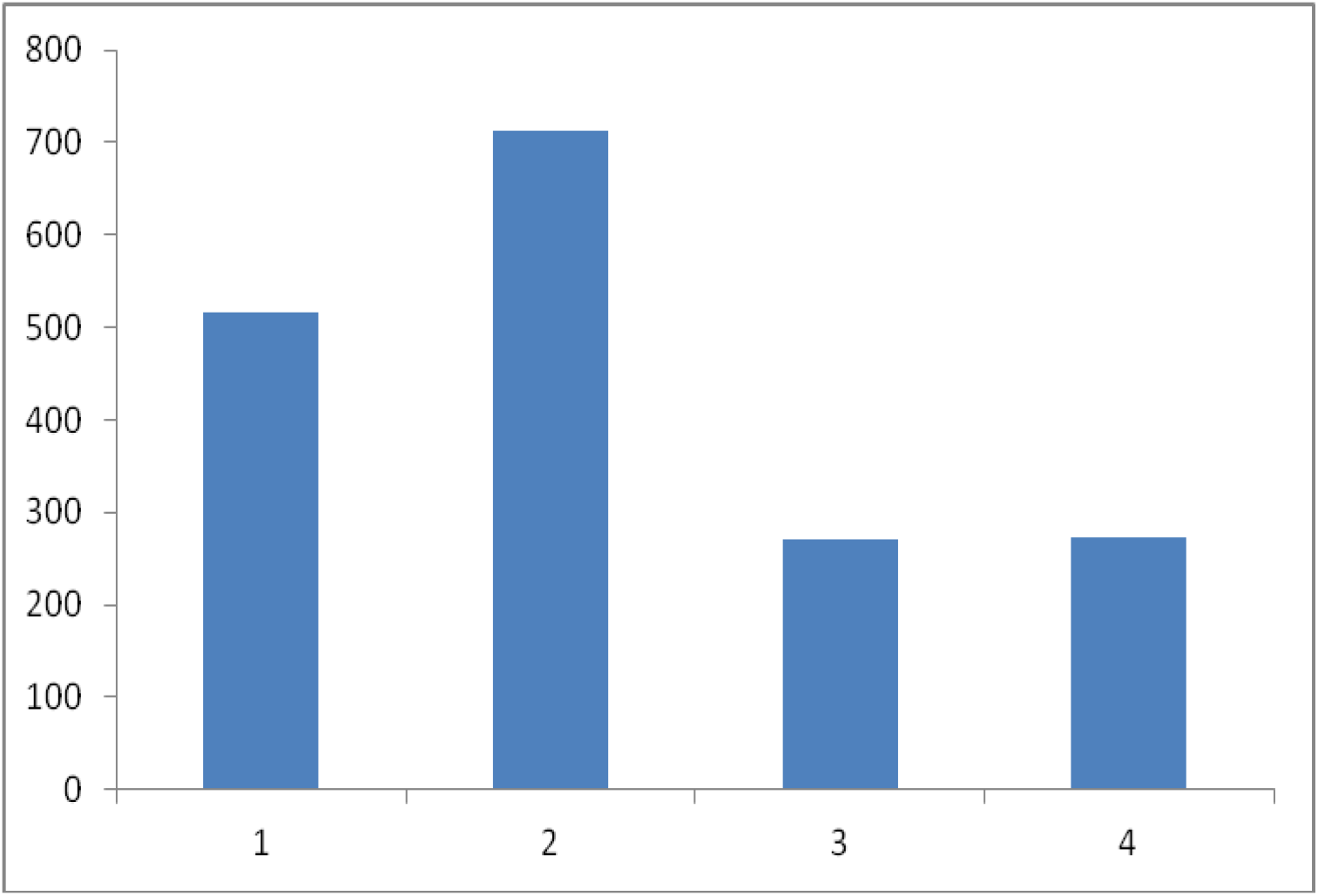
Total protein expression profile of soot and H2o2. Histograms represent soot induced total protein in the human lung epithelial cells. 1=control, 2= H_2_O_2,_ 3= H_2_O_2_ + soot 250 µg/ml, 4=soot alone 250 µg/ml

#### 4. Effect of autophagy inhibitor 3-methyl adenine (3-MA) on the DEP induced cell proliferation of A549 cells

Further to understand the mechanism of autophagy in response to DEP, we have tested the effects of autophagy inhibiter 3-methyl adenine (3-MA) on A549 cells. Different concentrations (250 µg/ml, 500, µg/ml and 1000 µg/ml) of DEP were exposed to lung A549 cells for a period of 24 hours in the presence and absence of 3MA. As seen in the figure 3-MA significantly decreased DEP induced cell proliferation of A549 cells in dose dependent manner.

**Supplementary figure 4:**
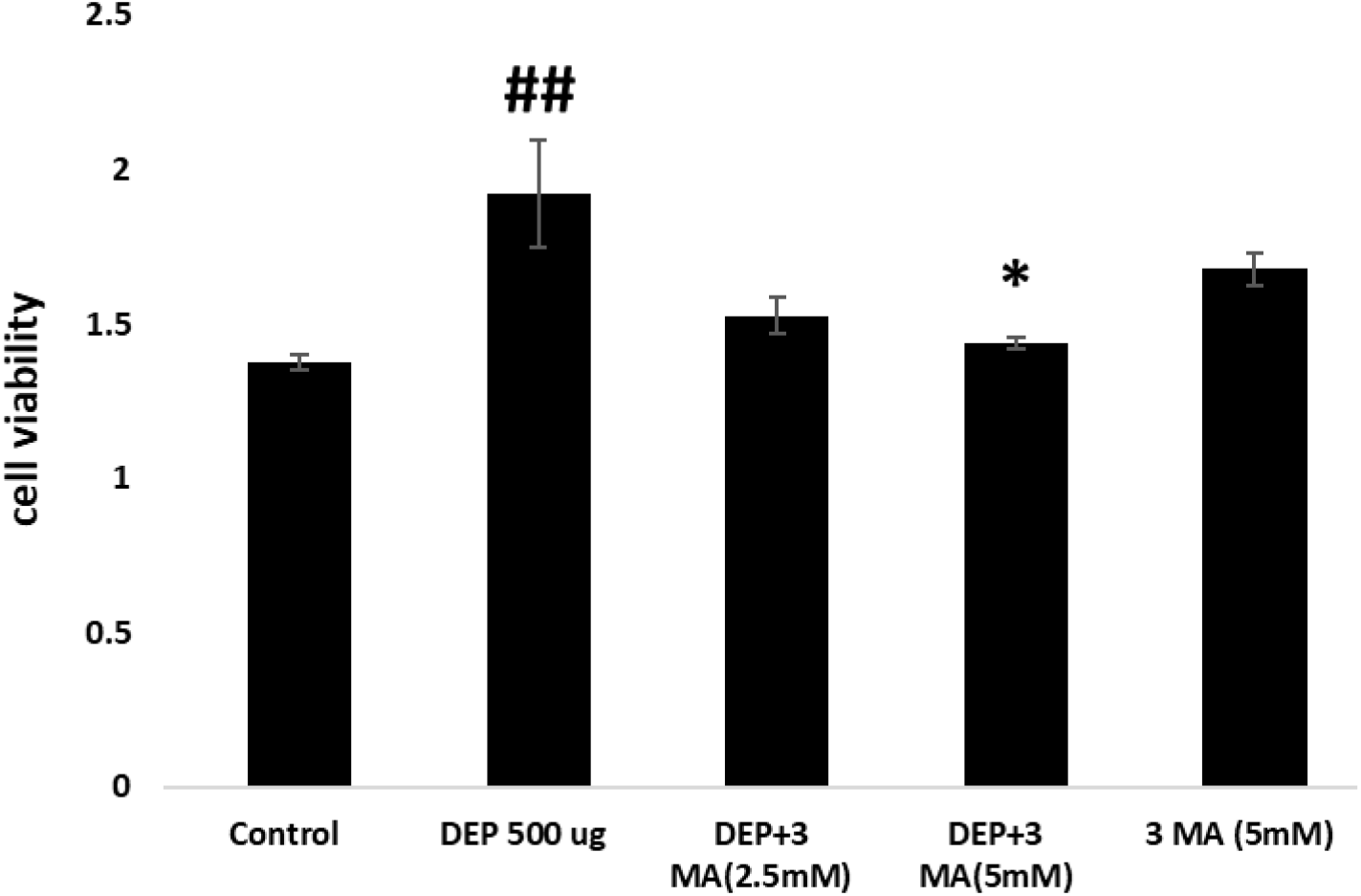
Effect of 3-MA on the DEP induced cell proliferation of A549 cells. Histogram represent relative cell proliferation of A549 cells expressed as Mean ± SE. # #p<0.01 compared with control, and * p<0.05 compared with DEP treated group.

